# Deformability based sorting of stored red blood cells reveals donor-dependent aging curves

**DOI:** 10.1101/818765

**Authors:** Emel Islamzada, Kerryn Matthews, Quan Guo, Aline T. Santoso, Simon P. Duffy, Mark D. Scott, Hongshen Ma

**Author notes:** Correspondence should be addressed to Hongshen Ma.

## Abstract

A fundamental challenge in the transfusion of red blood cells (RBCs) is that a subset of donated RBC units may not provide optimal benefit to transfusion recipients. This variability stems from the inherent ability of donor RBCs to withstand the physical and chemical insults of cold storage, which ultimately dictate their survival in circulation. The loss of RBC deformability during cold storage is well-established and has been identified as a potential biomarker for the quality of donated RBCs. While RBC deformability has traditionally been indirectly inferred from rheological characteristics of the bulk suspension, there has been considerable interest in directly measuring the deformation of RBCs. Microfluidic technologies have enabled single cell measurement of RBC deformation but have not been able to consistently distinguish differences between RBCs between healthy donors. Using the microfluidic ratchet mechanism, we developed a method to sensitively and consistently analyze RBC deformability. We found that the aging curve of RBC deformability varies significantly across donors, but is consistent for each donor over multiple donations. Specifically, certain donors seem capable of providing RBCs that maintain their deformability during two weeks of cold storage in standard test tubes. The ability to distinguish between RBC units with different storage potential could provide a valuable opportunity to identify donors capable of providing RBCs that maintain their integrity, in order to reserve these units for sensitive transfusion recipients.

## Introduction

Donated red blood cells (RBCs) are a central component of critical patient care with >85 million units transfused globally per year^1^. A fundamental challenge in the transfusion of donated RBCs is that not all donated RBC units can confer the same benefits to recipients^2,3^. Specifically, transfusion specialists have long been aware that certain RBC units are able to circulate for long periods of time in recipients to maintain hemostasis, while other units are rapidly cleared, leading to the need for repeat transfusions^4–8^. This situation is highly undesirable for chronic transfusion recipients because of potential adverse effects, such as iron-overload, transfusion related acute lung injury (TRALI), hypervolemia, and increased risk of infections^9–12^. Consequently, there is a strong impetus to establish robust biomarkers to identify donors that could provide long circulating RBC units in order to match them to the clinical needs of the patient^13,14^.

An essential characteristic of RBCs is their deformability, which enables these cells to transit through the microvasculature. The loss of deformability causes RBCs to be retained in the narrow inter-endothelial clefts of the spleen^15^, where they are subsequently removed from circulation^16^. During cold storage, it is well-established that donated RBCs lose their deformability as a part of the storage lesion^13,17–21^. While the storage lesion can contribute to a host of cellular changes, including depletion of 2,3-DPG^22^, cytoskeletal remodeling^23^, lipid oxidation^24,25^, and membrane vesiculation^26,27^, these effects independently as well as cumulatively contribute to the irreversible depletion of RBC deformability. Consequently, the deformability of RBC has long been considered as a potential biomarker for the quality of donated RBCs.

Existing methods for measuring RBC deformability include flow-based methods and deformation-based methods. Flow-based methods, including ektacytometry^28,29^ and related microfluidic approaches^30–32^, use shear stress generated in fluid flow to stretch RBCs in order to measure their elongation via diffraction or high-speed imaging. These approaches measure the rheological properties of RBCs during high-speed flow rather than the deformation of individual RBCs into the splenic microvasculature, which is the primary mechanism for RBC clearance. Deformation based methods include the traditional methods for single cell manipulation and recent microfluidic devices. Traditional single cell manipulation, including micropipette aspiration^33,34^, optical tweezers^35^, and atomic force microscopy^36^, involve technically challenging experiments that require skilled personnel and highly specialized equipment. Microfluidic approaches for deformation based RBC deformability analysis include the measurement of capillary obstruction^37^, wedging in tapered constrictions^38,39^, transiting time through constrictions^40–43^, and transiting pressure through constrictions^17,44,45^. While these approaches greatly simplified RBC deformability analysis, the throughput of these processes are still relatively limited. Therefore, a frequent criticism is potential bias arising from the cell sampling process. Furthermore, microfluidic deformation approaches often rely on precise geometry of a small number of microscopy features, which can vary from device to device. Consequently, previous methods are typically used to assay the loss of RBC deformability due to pathologies or chemical treatment, but not to assess variability between healthy individuals.

Recently, we developed a microfluidic device to sort RBCs based on deformability using a matrix of asymmetrical constrictions that form microfluidic ratchets^46–48^. Key advantages of this device are the ability to apply consistent forces to each cell as it deforms across each constriction, as well as the ability to process a large number of cells^48^. These characteristics improve the consistency and throughput of single cell deformation measurements by averaging the measurement across large numbers of individual RBCs. Leveraging this capability, we show deformability based cell sorting using microfluidic ratchets provides a highly sensitive and consistent measurement of RBC deformability. Specifically, this measurement is capable of detecting distinct and temporally consistent differences between blood units from healthy donors, which we observed to exhibit donor-specific aging rates during cold storage.

The ability to detect differences in RBC deformability between healthy donors, as well as the loss of integrity during cold storage, provides a potential opportunity to identify donors who may provide RBCs that maintain their integrity during cold storage. This capability will be extremely important for chronic transfusion recipients that require long-circulating RBC units in order to reduce the number of transfusions. Reserving high-quality RBC units for these patients will improve outcomes and increase the overall blood supply.

## Results

### Microfluidic device design

The microfluidic ratchet device sorts RBCs based on deformability by deforming each cell through a series of tapered constrictions. (**Fig. 1A**). Due to the geometric asymmetry of the taper, the force required to deform cells through the constriction along the direction of taper is less than against the direction of taper^46^. Coupling this deformation asymmetry with an oscillatory flow creates a ratcheting effect that selectively transports cells based on their ability to squeeze through each microscopic constriction. Importantly, this oscillatory flow also minimizes the contact between cells and the filter microstructure to prevent clogging and fouling and to ensure that a consistent filtration force is applied to each cell^47^.

**Fig. 1.**
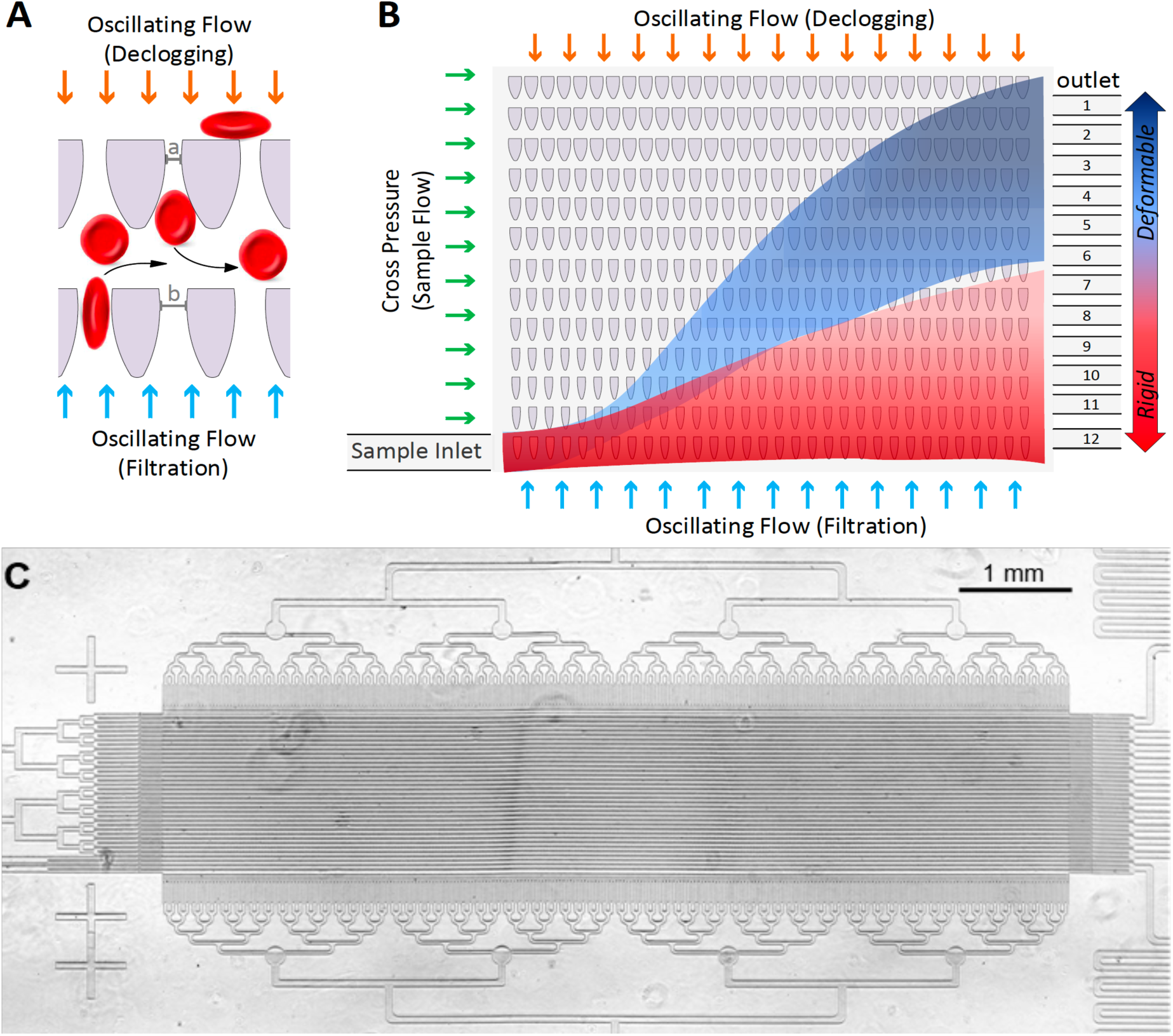
The microfluidic ratchet device for deformability-based cell sorting. (A) Tapered funnel constrictions enable unidirectional (upwards) transport of individual RBCs from oscillatory flow. Rows of constrictions decreasing in size (i.e. *a*<*b*) enable deformability based cell separation. (B) Deformability based cell sorting is performed using a matrix of funnel constrictions. The RBC sample is flowed into the lower left corner of the matrix and follows a diagonal trajectory resulting from vertical oscillatory flow and horizontal cross flow. RBCs travel diagonally through the matrix until reaching a limiting constriction size, which forces them to flow horizontally to a specific outlet. Deformable cells (blue shading) and rigid cells (red shading) follow distinct paths because of different limiting constrictions. (C) Micrograph of the microfluidic ratchet device.

To sort cells using this mechanism, a RBC sample is introduced at the bottom-left corner of a 2D array of micrometer-scale tapered constrictions. The openings of the constrictions are gradually decreased from the bottom row to the top row, forcing RBCs to deform as they transit through progressively smaller pores. The RBCs are transported through the array by a vertical oscillatory flow as well as a consistent horizontal cross flow, which combine to propel the cells in a zigzag diagonal path through the constriction matrix (**Fig. 1B**). As cells reach their limiting constriction that prevent their transport, they proceed horizontally along the terminal row until reaching a specific outlet (**Fig. 1C**). The microfluidic device used to sort RBCs from healthy donors is designed to sort RBCs into 12 outlets. The constriction matrix is housed in a 4 µm thick microchannel. The size of the constrictions ranges from 1.50 µm to 3.50 µm in steps of 0.25 µm for outlets 1-9, and then from 3.50 µm to 7.50 µm in greater steps for outlets 10-12 (**Table 1**). The 4 µm thick microchannel largely exclude leukocytes, debris and cell aggregates. However, if these artefacts enter the sorting area, they are sorted into the least deformable outlets, e.g. 10-12. As a result, we have not found evidence for nucleated cells or cell clusters in outlets 1-9. This device is capable of sorting ∼600 RBCs per minute. The distribution of cells after sorting could be determined by imaging the flow of cells into the outlets or by counting the cells in the outlet via microscopy.

**Table 1.** Outlet numbers and their corresponding constriction sizes.

| Outlet | 1 | 2 | 3 | 4 | 5 | 6 | 7 | 8 | 9 | 10 | 11 | 12 |
| --- | --- | --- | --- | --- | --- | --- | --- | --- | --- | --- | --- | --- |
| Size (μm) | 1.5 | 1.75 | 2 | 2.25 | 2.5 | 2.75 | 3 | 3.25 | 3.5 | 4.5 | 5.5 | 7.5 |

### RBC rigidity score

After deformability-based cell sorting, the distribution of RBCs in outlets 1-12 can be shown as a histogram, where the loss of RBC deformability results in a rightward shift in distribution (**Fig. 2A**). In order to effectively compare differences between samples, we can plot the result as a cumulative distribution, and then define a “rigidity score” as the outlet where the cumulative distribution function crosses 50%. A fractional value for the rigidity score can be obtained by linear interpolation between data points in the cumulative distribution function. For example, the deformable and rigid RBC samples have rigidity scores of 2.45 and 4.87 respectively (**Fig. 2B**).

**Fig. 2.**
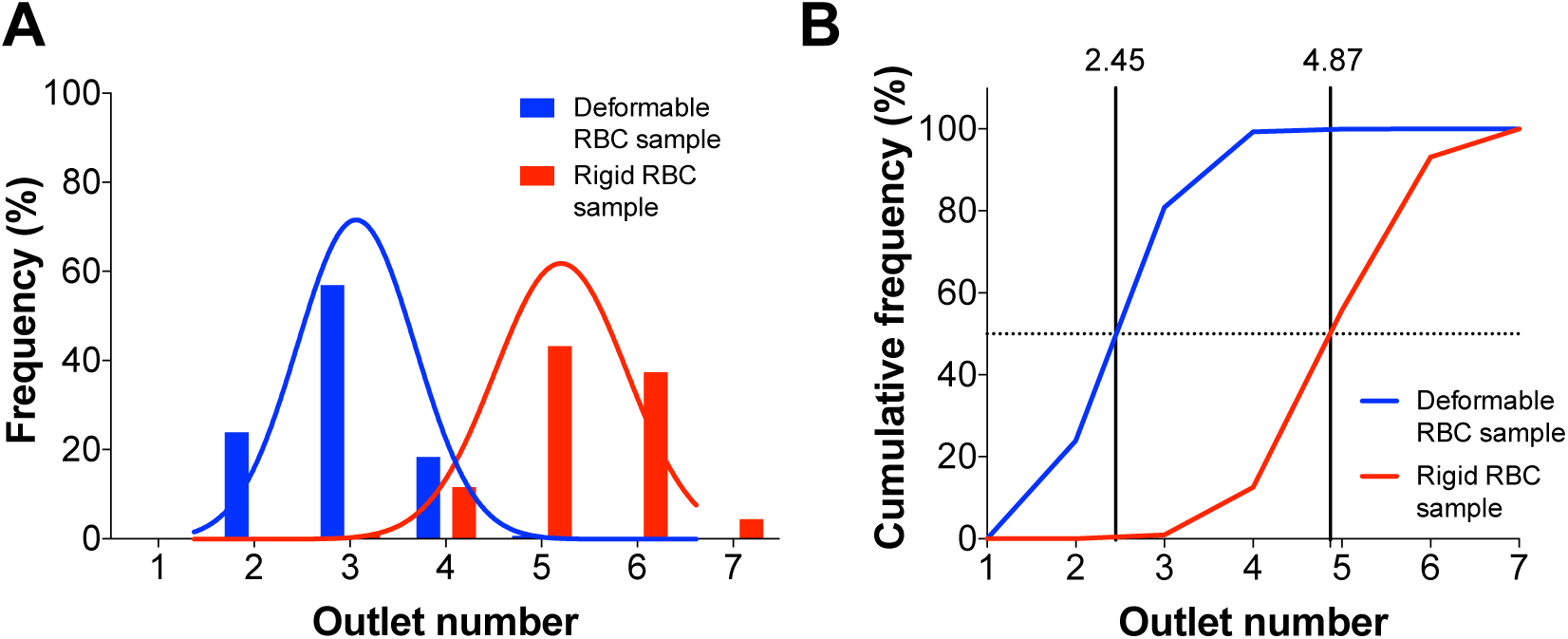
Analysis of deformability-based cell sorting data. (A) Distribution of RBCs after sorting shown as histograms for deformable (blue) and rigid (red) RBC samples. (B) Distribution of RBCs after sorting shown as cumulative distributions. The rigidity score (2.45 and 4.87 for the deformable and rigid RBC samples) is defined as the 50% cross-over point for the cumulative distribution.

### Process characterization and validation

We performed a series of studies to characterize the consistency and variability in our RBC sorting process. To characterize device-to-device variations, we used our process to sort 1.54 and 3.43 µm polystyrene calibration beads. The 1.53 µm beads were sorted into outlets 2 and 3, while the 3.43 µm beads were sorted into outlet 9. Similar results were obtained from 8 different devices, where the mean rigidity score for the 1.53 µm beads were 2.60 ± 0.24 (**Fig. 3A**). We then evaluated the consistency of the sorting process on human RBCs. We sorted RBCs from a single donor simultaneously using 5 different devices. The RBC distribution were also highly consistent, where the rigidity score for this sample was 2.63 ± 0.17 (**Fig. 3B**). Together, these data show that the device fabrication process is robust and that the experimental process yield consistent results.

**Fig. 3.**
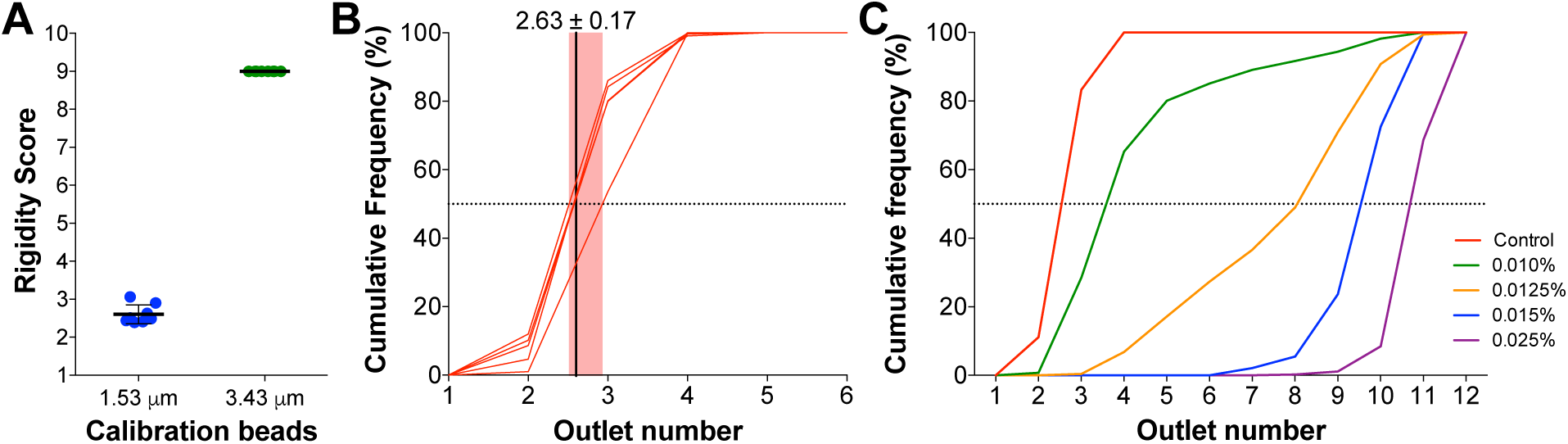
Characterization and Validation of the Cell Sorting Process. (A) Rigidity score from sorting 1.53 µm (8 repeats) and 3.43 µm (7 repeats) diameter polystyrene beads using devices from different manufacturing batches. Error bars indicate SD. (B) Cumulative distributions obtained from sorting the same RBC sample using 5 devices from different manufacturing batches shown with the mean rigidity score (solid line) and range (shading). (C) Cumulative distributions showing the loss of RBC deformability after mild glutaraldehyde fixation.

To establish that the rigidity score could discriminate between RBC populations with a range of cell deformability, we obtained fresh RBCs and applied mild glutaraldehyde (GTA) fixation in order to induce controlled rigidification of the cells. Following 30 minute GTA fixation, a dose-dependent rightward shift could be observed in the distribution of RBCs within the device outlets, following sorting (**Fig. 3C**). Compared to the control (rigidity 2.63 ±0.17), GTA-fixed cells displayed rigidity scores of 3.6 (0.010% GTA), 8.0 (0.0125% GTA), 9.5 (0.015% GTA) and 10.7 (0.025% GTA).

### Donor-based variability in RBC deformability

Having established sensitivity and consistency, we used our cell sorting approach to characterize donor-based variability in RBC deformability at the time of collection with 8 donors. Fresh RBCs are typically sorted into outlets 2-3 with some variability in their distribution. The 8 donors had a mean rigidity score 2.90 ± 0.4, ranging between 2.36 (donor 1) and 3.59 (donor 5) (**Fig. 4A**). To establish whether donor deformability profiles remained consistent over time, RBCs were collected from donors 1 and 5 on two more occasions, with at least 6 weeks between each sampling (**Fig. 4B**). Interestingly, the measured rigidity of RBCs for each donor was highly consistent for samples collected at each distinct occasion. Specifically, the rigidity score for donor 1 ranged from 2.36 to 2.57, while Donor 5 ranged from 3.19 to 3.59.

**Fig. 4.**
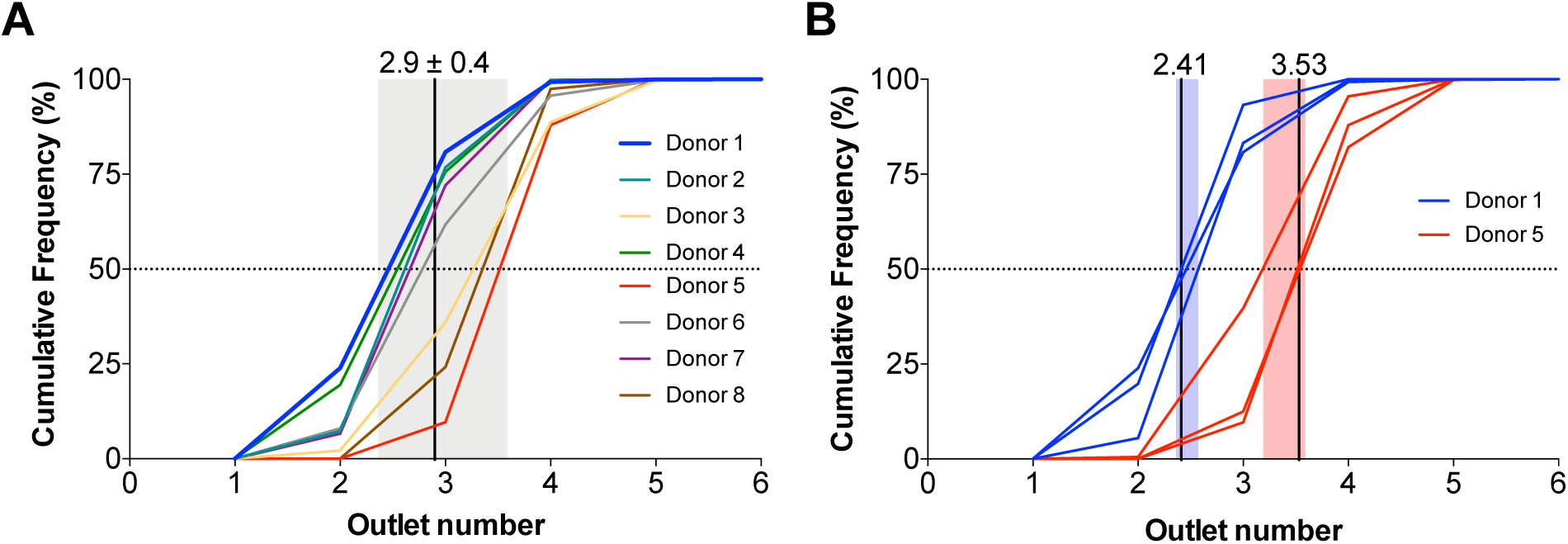
Inter-and Intra-donor variability of fresh RBC deformability. (A) Cumulative distributions from deformability based sorting of fresh RBC samples from 8 healthy donors. Mean rigidity score is shown as solid line and SD shown as shading. (B) Cumulative distributions of 3 repeat RBC samples from donors 1 (blue) and 5 (red) taken at least 6 weeks apart.

### Donor-based variability in RBC deformability aging curves

To establish whether cell sorting using the microfluidic ratchet device could provide a robust method to measure the degradation of RBCs during cold storage, we measured the loss of deformability associated with the storage of RBCs in a standard plastic tube, which provide a more consistent model of accelerated aging over 14 days compared to a blood bag^49^. Donated blood samples from all 8 donors were stored in standard 15 ml plastic test tubes (Corning) for 14 days at 4°C. Deformability based cell sorting was performed on day 0, 3, 7, and 14. Interestingly, while all samples showed some loss of RBC deformability during cold storage (mean rigidity score change = +1.0), there was significant variability between donors (**Fig. 5**). Specifically, donor 1 showed the most significant loss of deformability during cold storage (+2.51 in rigidity score), even though this donor had the most deformable RBCs at the point of collection (rigidity score = 2.36). Whereas RBCs from donor 5 and donor 8 were remarkably stable, where the rigidity score is essentially unchanged (−0.04 and +0.25 for donor 5 and 8 respectively, **Fig. 6**).

**Fig. 5.**
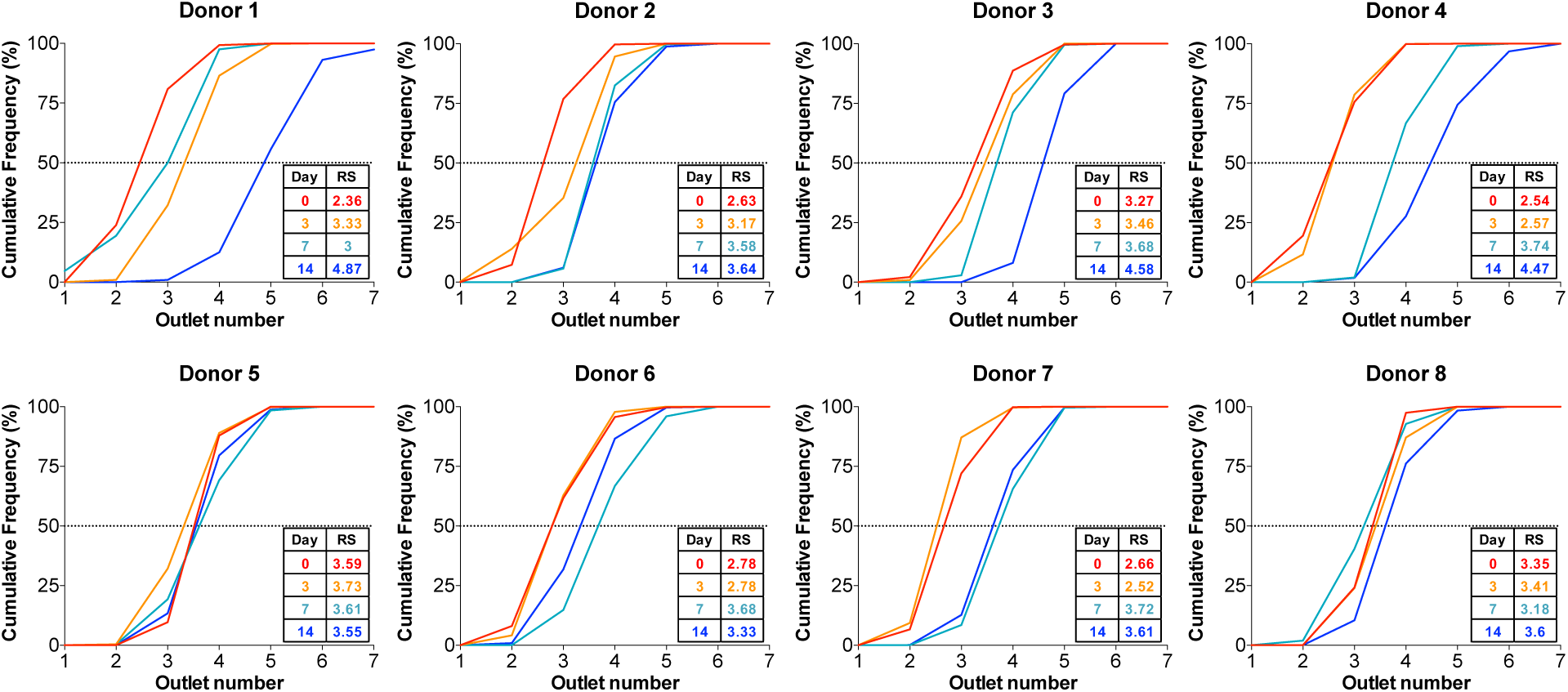
RBC deformability aging curves. Cumulative distributions curves from deformability-based sorting of RBCs obtained from 8 donors after 0, 3, 7, and 14 days of accelerated aging. Rigidity scores (RS) are listed in the inset tables. Significant loss of RBC deformability is observed from most donors after 14 days with the notable exception of donors 5 and 8, which demonstrate little change after 14 days.

**Fig. 6.**
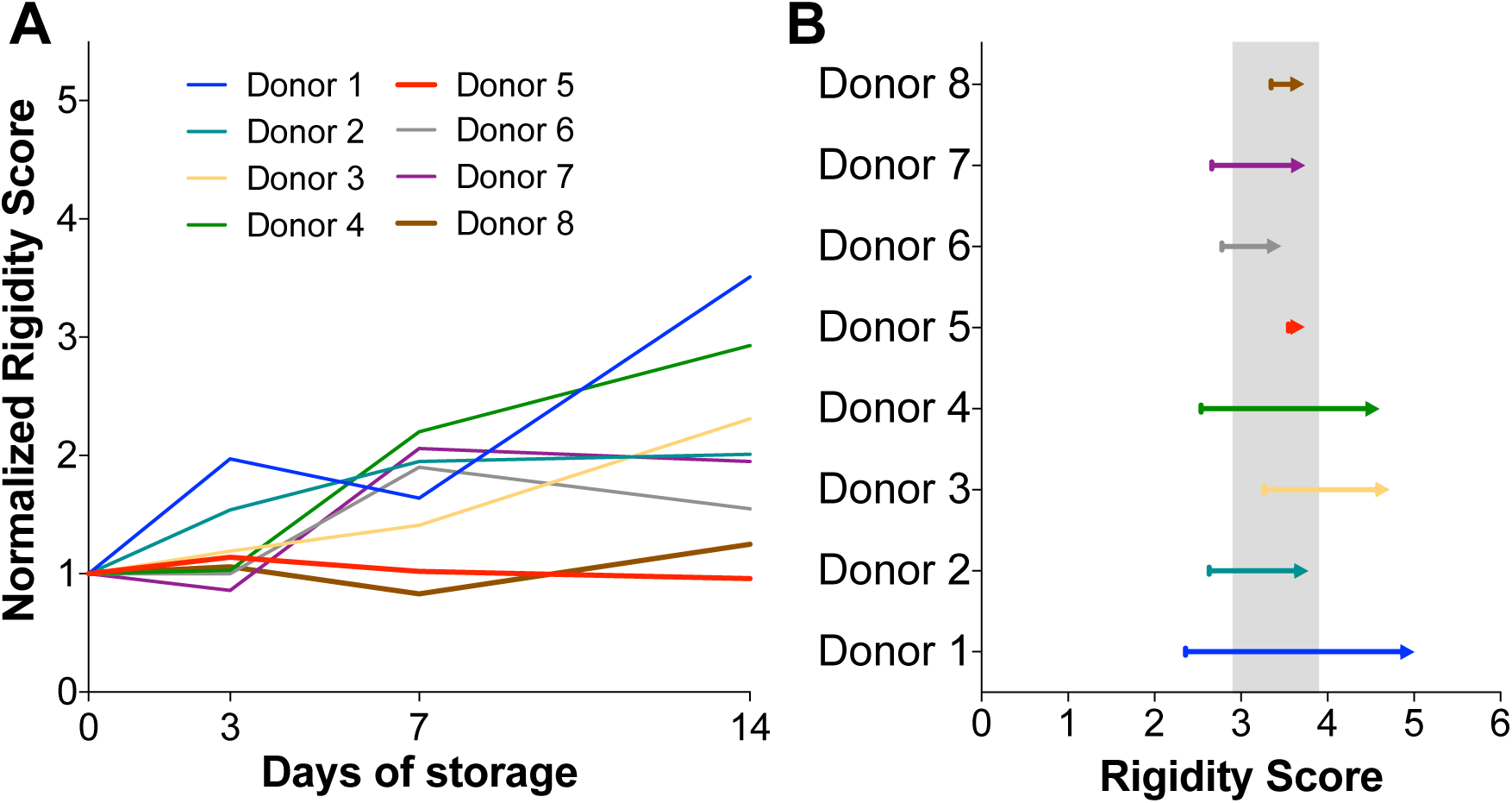
Summary of RBC deformability aging curves. (A) Normalized rigidity score from donors 1-8 over 14 days of accelerated aging. (B) Change in rigidity score from day 0 to day 14. The shading indicates the mean change for all 8 donors.

Having observed that fresh RBC deformability was consistent for each donor, we investigated whether the RBC aging curve was also donor-specific when blood was collected on different occasions. RBC samples were collected from donors 1 and 5 at 6-week intervals and RBC deformability profiles were assessed over the course of storage to establish the RBC aging curve. Consistent with our observation that RBC rigidity was temporally consistent for each donor, we also observed that the aging curve was remarkably consistent when sampled on different occasions (**Fig. 7**). Donor 1 consistently showed an accelerated RBC aging curve, while donor 5 consistently provided RBCs that retain their integrity during cold storage.

**Fig. 7.**
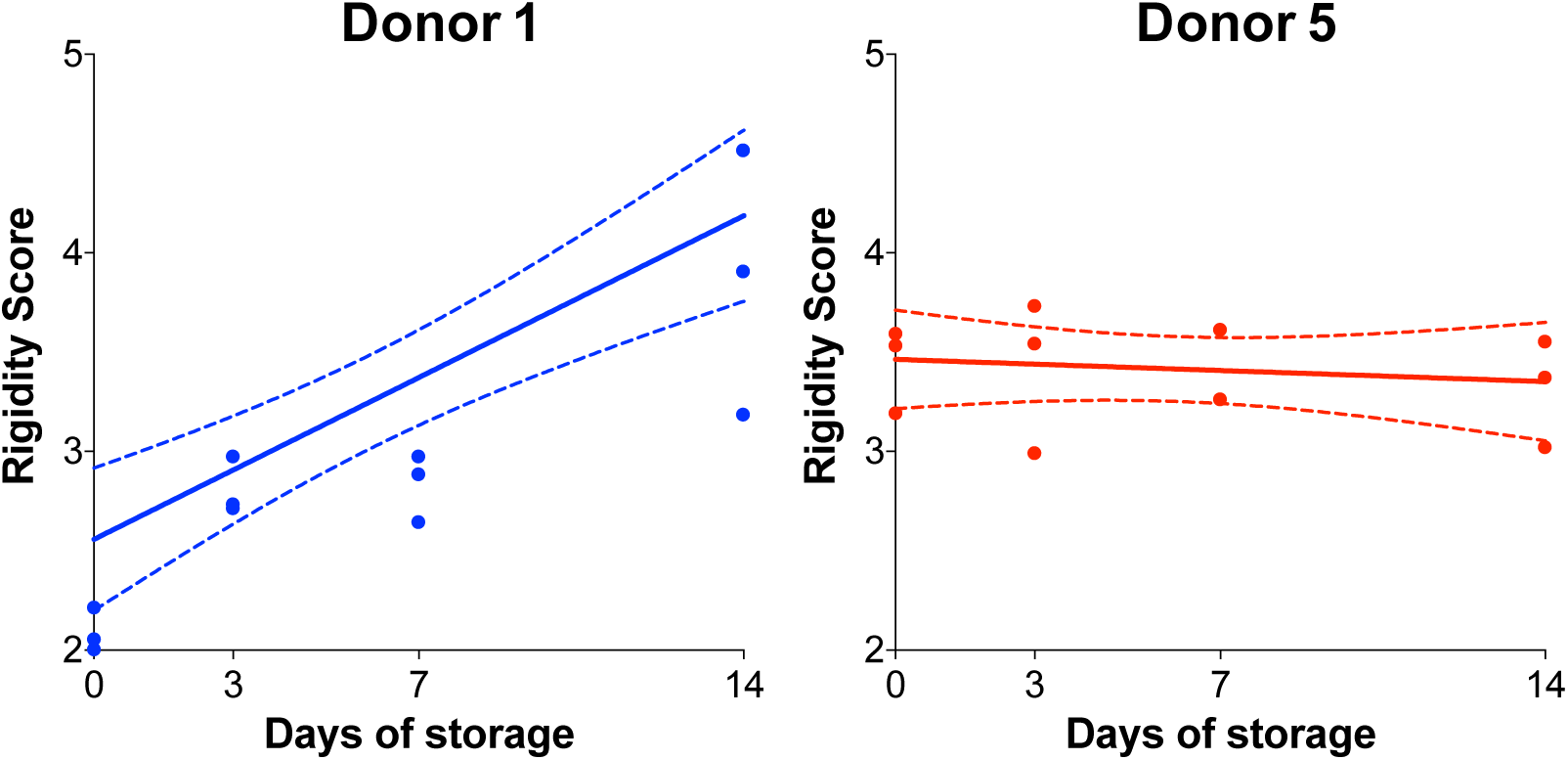
Intra-donor variability of RBC deformability aging profiles. Rigidity scores (dots) over 14 days of accelerated aging for 3 RBC samples obtained at least 6 apart from donors 1 and 5. Linear regression analysis (solid lines) showed that donor 1 (blue) had consistent loss of deformability during storage (R^2^=0.768, p=0.0002), while donor 5 (red) showed no differences between Day 0 and Day 14 of storage (R^2^=0.032, p=0.5799).

### Hematological parameters do not affect RBC sorting

We also measured standard hematological indices during storage, including RBC mean corpuscular volume (MCV), red blood cell distribution width (RDW-CV), mean cell hemoglobin (MCH), and mean corpuscular hemoglobin concentration (MCHC), using the SYSMEX® system. There was a trend of increasing RBC MCV and MCHC during storage in the plastic tubes over the 14 days, while the RDW-CV and MCH remained largely unchanged (Table S1). However, these parameters all remained within the normal range during storage. Furthermore, linear regression analysis of these parameters and RBC rigidity score, did not show any significant correlations (**Fig. S1**).

## Discussion

In this study we employed the microfluidic ratchet mechanism to sort RBCs from healthy donors based on deformability. After sorting, the RBCs in each outlet were counted and the distribution of cells was used to establish a rigidity score for the RBC population. Using this approach, we found that the baseline RBC deformability varies significantly between donors, but is relatively consistent for the same donor over multiple donations. Importantly, we found the RBC aging curve during cold storage, as defined by the loss of RBC deformability, is also heterogeneous, but consistent for each donor. These results collectively suggest RBC deformability measurements performed by sorting using the microfluidic ratchet could be used to identify donors capable of providing RBCs that maintain their integrity during cold storage.

RBC deformability is a key part of the mechanism for splenic clearance of RBCs^50,51^, and has therefore been frequently proposed as a potential biomarker for estimating the clearance time of transfused RBCs^52,53^. Traditional methods, such as ektacytometry^28,29,54,55^, test the rheological properties of RBC derived from bulk flow, which does not correlate well with circulatory clearance time^56,57^. Microfluidic technologies present the potential to improve upon this measurement process by using microfabricated structures to approximate the deformation mechanics of the splenic microvasculature^58–60^. However, key limitations of existing microfluidic devices is the throughput of the measurement process, as well as sample-to-sample consistency. Using the microfluidic ratchet mechanism to perform continuous deformability-based cell sorting, we greatly increased measurement throughput. Importantly, by significantly increasing redundancy of the microstructure, we showed our process to be highly consistent and repeatable. The ability to perform consistent high-throughput measurement of RBC deformability will permit the study of the relationship between RBC deformability and circulation time, using approaches such as the recovery rate of ^51^Cr- or biotin-labeled RBCs in transfused animal models^61–64^.

RBC deformability measured using the microfluidic ratchet revealed significant inter-donor variability of freshly collected RBCs as well as inter-donor variability in RBC rigidification during storage. Interestingly, while previous efforts have reported storage-mediated mechanical deterioration of RBCs, deformability of fresh RBCs has largely been reported to be invariant^56,57^. Together, inter-donor variability in RBC deformability of both fresh and stored RBCs supports epidemiological evidence that RBC transfusion and storage efficacy is donor-dependent^65–69^. While the data obtained in this study are too limited to derive associations between RBC deformability and transfusion efficacy, the observation that individual donors exhibit distinct and consistent RBC deformability before and during storage enables future studies to correlate donor RBC deformability and aging curves, as well as genetic, biochemical, and hematological rheological parameters, with clinical outcomes and the frequency of adverse events posttransfusion. An association between donor RBC deformability and transfusion efficacy could lead to the stratification of donors based on transfusion and RBC storage potential.

The identification of donors that can consistently provide high quality RBCs could have a profound impact on transfusion medicine. RBC units with a longer predicted circulation time could be matched to chronic transfusion recipients in order to reduce the frequency of transfusions and thereby, reduce transfusion-related morbidities, including iron-overload, hypervolemia, transfusion related acute lung injury (TRALI), as well as potential infection^9–11^. Furthermore, the ability to predict the circulation time of RBCs based on cell deformability could provide new methods of quality control in order to improve quality and reduce excessive outdating. In sum, the ability to match donated RBC units with the most appropriate recipient will translate to improved outcomes for sensitive recipients and optimal usage of this limited resource.

## Methods

### RBC sample collection, storage, and preparation

Only donors self-identified as healthy and between the ages of 18 and 70 were included in this study. Following informed consent and in accordance with University of British Columbia Research Ethics Board guidelines (UBC REB H19-01121), whole blood was collected from healthy donors via venipuncture into sodium citrate tubes (BD Vacutainer). The blood samples were leuko-reduced by first centrifuging at 3180 g for 10 minutes to decant plasma. The packed cells were then centrifuged at 677 g for 30 minutes with no brakes to remove the leukocytes. Each 1 mL of packed RBC was supplemented with 0.80 mL of saline-adenine-glucose-mannitol storage solution (SAGM) and 0.56 mL of plasma, and stored in a 15 mL plastic tube (Corning) upright at 4°C. On days 0, 3, 7, and 14, an aliquot of the sample was taken for RBC deformability measurement with the Microfluidic Ratchet device. The RBC pellet was resuspended in Hanks Balanced Salt Solution (HBSS, Gibco) and 0.2% Pluronic solution (F127, MilliporeSigma) at 1% hematocrit for infusion into the microfluidic ratchet device. A second unwashed sample aliquot was used to measure standard hematological indices, such as RBC mean corpuscular volume (MCV), red blood cell distribution width (RDW-CV), mean cell hemoglobin (MCH), and mean corpuscular hemoglobin concentration (MCHC), using the SYSMEX® system.

### Microfluidic ratchet device manufacture

The microfluidic ratchet devices were prepared as described previously^70,71^. Briefly, the master device mold was created by photolithographic microfabrication. This mold was then used to create a secondary polyurethane mold, fabricated with Smooth-Cast urethane resin (Smooth-Cast ONYX SLOW, Smooth-On) as described previously^72^. Single-use microfluidic devices were molded from the secondary master using PDMS silicone (Sylgard-184, Ellsworth Adhesives) mixed at a 10:1 ratio with the curing agent (Sylgard-184, Ellsworth Adhesives). The PDMS devices were cured for two hours at 65°C. After removing the PDMS devices from the molds, the sample inlets and outlets were manually punched with 0.5 mm and 3.0 mm hole punches, respectively. (Technical Innovations). The bottom surface of the microfluidic devices was created using a thin layer of PDMS silicone (RTV 615, Momentive Performance Materials LLC). This thin layer of PDMS silicone was formed by spin coating the uncured PDMS on a 100 mm silicon wafer at 1500 RPM for 1 minute and then curing at 65°C for two hours. The Sylgard-184 PDMS microstructure was then bonded to the RTV 615 thin PDMS layer using air plasma (Model PDC-001, Harrick Plasma). The composite structure was then bonded to a 75 x 50 mm glass slide (Corning) substrate using air plasma.

### Microfluidic device operation

The microfluidic ratchet device is operated using a horizontal crossflow and oscillatory (forward/reverse) pressure system. Prior to loading the RBC sample, the device is primed by infusion with HBSS with 0.2% Pluronic-F127 solution, through the forward and oscillatory pressure system inlets at high pressure (200-250 mBar) for 15 min. The RBC sample for each assay is suspended at 1% hematocrit and then infused into the constriction matrix using a pressure of 40-45 mBar through the sample inlet port. The sample is then flowed through the constriction matrix using a horizontal crossflow pressure of 55-60 mBar, and oscillatory pressures of 175 mBar forward and 162 mBar reverse (**Fig. 1**). The forward oscillatory pressure is applied for 4 seconds, while the reverse oscillatory pressure is applied for 1 second. The crossflow pressure is adjusted visually within the pressure range to ensure the equal distribution of cells within the sample inlet as well as a consistent application of upward pressure to each cell. The throughput of the sorting process is approximately 600 cells per minute. From the constriction matrix, the RBC sample is flowed into 12 outlets. The distribution of cells after sorting can be counted via video analysis of the microchannels leading to the outlets, or by imaging the outlets. The sorted RBCs could also be collected by pipetting.

### Microbead sorting experiments

Polystyrene microbeads were sorted using the microfluidic ratchet device in order to establish the consistency across different microfluidic devices. Polystyrene beads with diameters of 1.53 µm and 3.42 µm (Cat # 17133, Polysciences Inc) were selected to coincide with approximate deformed sizes of deformable and rigid RBCs. Beads were vortexed prior to opening, and 25 µl of the bead solution was suspended in 650 µl of HBSS with 0.2% Pluronic F127 and 0.2% TWEEN-20 (MilliporeSigma) solution to prevent aggregation, to a final concentration of 0.1 % beads in buffer. Each bead suspension was then sorted for at least 20 minutes. The number of beads in each outlet fraction was quantified using the TrackMate particle analysis plug-in with Image J^73^.

### RBC sorting experiments

Blood from a healthy donor was collected in a sodium citrate tube (BD Vacutainer), and RBCs were isolated as previously described above. The sample was then resuspended in HBSS and 0.2% Pluronic F127 solution to 1% hematocrit for further sorting with the microfluidic ratchet device. The sample from the same donor and suspension was run on 5 separate devices from different production batches. Sorted fractions were quantified using TrackMate particle analysis plug-in with Image J, as previously described.

### Glutaraldehyde fixation sorting experiments

In order to establish the sensitivity of the microfluidic ratchet device in measuring small changes in the deformability, RBCs from a healthy donor were collected in sodium citrate tubes (BD Vacutainer) and leuko-reduced as described above. The isolated RBCs were then washed three times and suspended at 10% hematocrit in HBSS with 0.02% Pluronic F127. These suspensions were fixed with either 0% (PBS control), 0.010%, 0.0125%, 0.015%, or 0.025% fresh glutaraldehyde (Cat # G5882, MilliporeSigma) for 30 minutes at room temperature. After fixation, the suspensions were washed three times in HBSS with 0.2% Pluronic F127 and then suspended at 1% Hct for the deformability experiment. The number of cells in each sorted fraction was quantified with the TrackMate particle analysis plug-in with Image J.

### Statistical Analysis

Deformability and rigidity score data were analysed with GraphPad Prism (v6.0h) software. Mean with standard deviation (SD) were plotted unless otherwise stated. Linear regression analysis with 95% confidence intervals were used to determine correlations between data sets.

## Acknowledgments

This work was supported by a CIHR/NSERC Collaborative Health Research Projects grant (414861), a CIHR Project Grant (362500), as well as an NSERC Discovery Grant (2015-06541). E.I. acknowledges funding from the Canadian Blood Services Graduate Fellowship. K.M. acknowledges funding from MITACS Accelerate program (IT09621). H.M. acknowledges funding from the CIHR New Investigator Salary Award program (322375).

## Authorship Contributions

H.M. conceived the idea and supervised the work. E.I., K.M., and Q.G. performed the experimental work. E.I., A.T.S., and Q.G. designed and manufactured the microfluidic devices used in the study. All authors wrote the manuscript.

## References

1. Takei T, Amin NA, Schmid G, Dhingra-Kumar N, Rugg D. Progress in global blood safety for HIV. J. Acquir. Immune Defic. Syndr. 2009;52 Suppl 2:S127–131.

2. Hess JR. Conventional blood banking and blood component storage regulation: opportunities for improvement. Blood Transfus. 2010;8 Suppl 3:s9–15.

3. Garraud O, Tissot J-D. Blood and Blood Components: From Similarities to Differences. Front Med (Lausanne). 2018;5:84.

4. Franco RS. Measurement of red cell lifespan and aging. Transfus Med Hemother. 2012;39(5):302–307.

5. Cohen RM, Franco RS, Khera PK, et al. Red cell life span heterogeneity in hematologically normal people is sufficient to alter HbA1c. Blood. 2008;112(10):4284–4291.

6. Carson JL. Red Blood Cell Transfusion: A Clinical Practice Guideline From the AABB*. Annals of Internal Medicine. 2012;157(1):49.

7. Tzounakas VL, Kriebardis AG, Papassideri IS, Antonelou MH. Donor-variation effect on red blood cell storage lesion: A close relationship emerges. PROTEOMICS - Clinical Applications. 2016;10(8):791–804.

8. Tzounakas VL, Georgatzakou HT, Kriebardis AG, et al. Donor variation effect on red blood cell storage lesion: a multivariable, yet consistent, story. Transfusion. 2016;56(6):1274–1286.

9. Wagner SJ. Transfusion-transmitted bacterial infection: risks, sources and interventions. Vox Sang. 2004;86(3):157–163.

10. Hendrickson JE, Hillyer CD. Noninfectious serious hazards of transfusion. Anesth. Analg. 2009;108(3):759–769.

11. Bux J. Transfusion-related acute lung injury (TRALI): a serious adverse event of blood transfusion. Vox Sanguinis. 2005;89(1):1–10.

12. Ng MSY, Ng ASY, Chan J, Tung J-P, Fraser JF. Effects of packed red blood cell storage duration on post-transfusion clinical outcomes: a meta-analysis and systematic review. Intensive Care Med. 2015;41(12):2087–2097.

13. D’Alessandro A, Kriebardis AG, Rinalducci S, et al. An update on red blood cell storage lesions, as gleaned through biochemistry and omics technologies. Transfusion. 2015;55(1):205–219.

14. Tzounakas VL, Kriebardis AG, Georgatzakou HT, et al. Glucose 6-phosphate dehydrogenase deficient subjects may be better “storers” than donors of red blood cells. Free Radic. Biol. Med. 2016;96:152–165.

15. Mebius RE, Kraal G. Structure and function of the spleen. Nat. Rev. Immunol. 2005;5(8):606–616.

16. Burger P, Hilarius-Stokman P, De Korte D, Van Den Berg T, Van Bruggen R. CD47 functions as a molecular switch for erythrocyte phagocytosis. Blood. 2012;119(23):5512–5521.

17. Matthews K, Myrand-Lapierre M-E, Ang RR, et al. Microfluidic deformability analysis of the red cell storage lesion. Journal of Biomechanics. 2015;48(15):4065–4072.

18. Tinmouth A, Fergusson D, Yee IC, Hébert PC, ABLE Investigators and the Canadian Critical Care Trials Group. Clinical consequences of red cell storage in the critically ill. Transfusion. 2006;46(11):2014–2027.

19. Pavenski K, Saidenberg E, Lavoie M, Tokessy M, Branch D. Red Blood Cell Storage Lesions and Related Transfusion Issues: A Canadian Blood Services Research and Development Symposium. Transfusion Medicine Reviews. 2011;26(1):68–84.

20. Zallen G, Moore EE, Blackwell J, et al. Age of transfused blood is an independent risk factor for postinjury multiple organ failure. Am J Surg. 1999;178(6):570–572.

21. Bennett-Guerrero E, Veldman TH, Doctor A, et al. Evolution of adverse changes in stored RBCs. Proc. Natl. Acad. Sci. U.S.A. 2007;104(43):17063–17068.

22. Li Y, Xiong Y, Wang R, Tang F, Wang X. Blood banking-induced alteration of red blood cell oxygen release ability. Blood Transfus. 2016;14(3):238–244.

23. Rael LT, Bar-Or R, Ambruso DR, et al. The effect of storage on the accumulation of oxidative biomarkers in donated packed red blood cells. J Trauma. 2009;66(1):76–81.

24. Racek J, Herynková R, Holecek V, Jerábek Z, Sláma V. Influence of antioxidants on the quality of stored blood. Vox Sang. 1997;72(1):16–19.

25. Wither M, Dzieciatkowska M, Nemkov T, et al. Hemoglobin oxidation at functional amino acid residues during routine storage of red blood cells. Transfusion. 2016;56(2):421–426.

26. Bosman GJCGM, Lasonder E, Luten M, et al. The proteome of red cell membranes and vesicles during storage in blood bank conditions. Transfusion. 2008;48(5):827–835.

27. Jy W, Ricci M, Shariatmadar S, et al. Microparticles in Stored RBC as Potential Mediators of Transfusion Complications. Transfusion. 2011;51(4):886–893.

28. Baskurt OK, Hardeman MR, Uyuklu M, et al. Comparison of three commercially available ektacytometers with different shearing geometries. Biorheology. 2009;46(3):251–64.

29. Streekstra GJ, Dobbe JGG, Hoekstra a G. Quantification of the fraction poorly deformable red blood cells using ektacytometry. Optics express. 2010;18(13):14173–82.

30. Forsyth AM, Wan JD, Ristenpart WD, Stone HA. The dynamic behavior of chemically “stiffened” red blood cells in microchannel flows. Microvasc Res. 2010;80(1):37–43.

31. Ahn KH, Lee SS, Yim Y, Lee SJ. Extensional flow-based assessment of red blood cell deformability using hyperbolic converging microchannel. Biomed Microdevices. 2009;11(5):1021–1027.

32. Katsumoto Y, Tatsumi K, Doi T, Nakabe K. Electrical classification of single red blood cell deformability in high-shear microchannel flows. Int J Heat Fluid Fl. 2010;31(6):985–995.

33. Hebbel RP, Leung a, Mohandas N. Oxidation-induced changes in microrheologic properties of the red blood cell membrane. Blood. 1990;76(5):1015–1020.

34. Evans E a, Waugh R, Melnik L. Elastic area compressibility modulus of red cell membrane. Biophysical journal. 1976;16(6):585–595.

35. Hénon S, Lenormand G, Richert a, Gallet F. A new determination of the shear modulus of the human erythrocyte membrane using optical tweezers. Biophysical journal. 1999;76(2):1145–1151.

36. Lekka M, Fornal M, Pyka-Fościak G, et al. Erythrocyte stiffness probed using atomic force microscope. Biorheology. 2005;42(4):307–317.

37. Shelby JP, White J, Ganesan K, Rathod PK, Chiu DT. A microfluidic model for single-cell capillary obstruction by Plasmodium falciparum infected erythrocytes. PNAS. 2003;100(25):14618–14622.

38. Herricks T, Antia M, Rathod P. Deformability limits of Plasmodium falciparum-infected red blood cells. Cellular Microbiology. 2009;11(9):1340–1353.

39. Gifford SC, Derganc J, Shevkoplyas SS, Yoshida T, Bitensky MW. A detailed study of time-dependent changes in human red blood cells: from reticulocyte maturation to erythrocyte senescence. Brit J Haematol. 2006;135(3):395–404.

40. Bow H, Pivkin I V, Diez-Silva M, et al. A microfabricated deformability-based flow cytometer with application to malaria. Lab on a chip. 2011;11(6):1065–73.

41. Zheng Y, Shojaei-Baghini E, Azad A, Wang C, Sun Y. High-throughput biophysical measurement of human red blood cells. Lab on a chip. 2012;12(14):2560–7.

42. Santoso AT, Deng X, Lee J-H, et al. Microfluidic cell-phoresis enabling high-throughput analysis of red blood cell deformability and biophysical screening of antimalarial drugs. Lab Chip. 2015;15(23):4451–4460.

43. Adamo A, Sharei A, Adamo L, et al. Microfluidics-based assessment of cell deformability. Analytical chemistry. 2012;84(15):6438–43.

44. Guo Q, Reiling SJ, Rohrbach P, Ma H. Microfluidic biomechanical assay for red blood cells parasitized by Plasmodium falciparum. Lab on a chip. 2012;12(6):1143–50.

45. Myrand-Lapierre M-E, Deng X, Ang RR, et al. Multiplexed fluidic plunger mechanism for the measurement of red blood cell deformability. Lab Chip. 2015;15(1):159–167.

46. Guo Q, McFaul SM, Ma H. Deterministic microfluidic ratchet based on the deformation of individual cells. Physical Review E. 2011;83(5):051910.

47. McFaul SM, Lin BK, Ma H. Cell separation based on size and deformability using microfluidic funnel ratchets. Lab Chip. 2012;12(13):2369–2376.

48. Park ES, Jin C, Guo Q, et al. Continuous Flow Deformability-Based Separation of Circulating Tumor Cells Using Microfluidic Ratchets. Small. 2016;12(14):1909–1919.

49. Lozano M, Cid J. DEHP plasticizer and blood bags: challenges ahead. ISBT Science Series. 2013;8(1):127–130.

50. Cluitmans J, Hardeman M, Dinkla S, Brock R, Bosman G. Red blood cell deformability during storage: towards functional proteomics and metabolomics in the Blood Bank. Blood Transfus. 2012;10 Suppl 2:s12–8.

51. Safeukui I, Buffet PA, Deplaine G, et al. Sensing of red blood cells with decreased membrane deformability by the human spleen. Blood Adv. 2018;2(20):2581–2587.

52. Koch CG, Duncan AI, Figueroa P, et al. Real Age: Red Blood Cell Aging During Storage. Ann. Thorac. Surg. 2019;107(3):973–980.

53. Bosch FH, Werre JM, Schipper L, et al. Determinants of Red-Blood-Cell Deformability in Relation to Cell Age. Eur J Haematol. 1994;52(1):35–41.

54. Bessis M, Mohandas N, Feo C. Automated ektacytometry: a new method of measuring red cell deformability and red cell indices. Blood cells. 1980;6(3):315–27.

55. Clark MR, Mohandas N, Shohet SB. Osmotic gradient ektacytometry: comprehensive characterization of red cell volume and surface maintenance. Blood. 1983;61(5):899–910.

56. Jani VP, Mailo S, Athar A, et al. Blood Quality Diagnostic Device Detects Storage Differences Between Donors. IEEE Trans Biomed Circuits Syst. 2017;11(6):1400–1405.

57. Tarasev M, Alfano K, Chakraborty S, et al. Similar donors-similar blood? Transfusion. 2014;54(3 Pt 2):933–941.

58. Sosa JM, Nielsen ND, Vignes SM, Chen TG, Shevkoplyas SS. The relationship between red blood cell deformability metrics and perfusion of an artificial microvascular network. Clin. Hemorheol. Microcirc. 2013;

59. Hou HW, Bhagat AAS, Chong AGL, et al. Deformability based cell margination--a simple microfluidic design for malaria-infected erythrocyte separation. Lab Chip. 2010;10(19):2605–2613.

60. Henry E, Holm SH, Zhang Z, et al. Sorting cells by their dynamical properties. Sci Rep. 2016;6(1):34375.

61. Mock D, Matthews N, Zhu S, et al. Red blood cell (RBC) survival determined in humans using RBCs labeled at multiple biotin densities. Transfusion. 2010;51(5):1047–1057.

62. Mock DM, Lankford GL, Widness JA, et al. Measurement of circulating red cell volume using biotin-labeled red cells: validation against 51Cr-labeled red cells. Transfusion. 1999;39(2):149–155.

63. Mollison PL. Further observations on the normal survival curve of 51 Cr-labelled red cells. Clin Sci. 1961;21:21–36.

64. Moroff G, Sohmer PR, Button LN. Proposed standardization of methods for determining the 24-hour survival of stored red cells. Transfusion. 1984;24(2):109–114.

65. Hess JR, Sparrow RL, Meer PF, et al. Red blood cell hemolysis during blood bank storage: using national quality management data to answer basic scientific questions. Transfusion. 2009;49(12):2599–2603.

66. Dern RJ, Gwinn RP, Wiorkowski JJ. Studies on the preservation of human blood. I. Variability in erythrocyte storage characteristics among healthy donors. J. Lab. Clin. Med. 1966;67(6):955–965.

67. Dumont LJ, AuBuchon JP. Evaluation of proposed FDA criteria for the evaluation of radiolabeled red cell recovery trials. Transfusion. 2008;48(6):1053–1060.

68. McAteer MJ, Dumont LJ, Cancelas J, et al. Multi-institutional randomized control study of haemolysis in stored red cell units prepared manually or by an automated system. Vox Sang. 2010;99(1):34–43.

69. Jordan A, Chen D, Yi Q-L, et al. Assessing the influence of component processing and donor characteristics on quality of red cell concentrates using quality control data. Vox Sang. 2016;111(1):8–15.

70. Guo Q, Duffy SP, Matthews K, et al. Deformability based sorting of red blood cells improves diagnostic sensitivity for malaria caused by Plasmodium falciparum. Lab Chip. 2016;16(4):645–654.

71. Guo Q, Duffy SP, Matthews K, Islamzada E, Ma H. Deformability based Cell Sorting using Microfluidic Ratchets Enabling Phenotypic Separation of Leukocytes Directly from Whole Blood. Scientific Reports. 2017;7(1):6627.

72. Desai SP, Freeman DM, Voldman J. Plastic masters-rigid templates for soft lithography. Lab Chip. 2009;9(11):1631–1637.

73. Abramoff M, Magalhaes P, Ram S. Image Processing with ImageJ. Biophotonics International. 2004;11(7):36–42.

